# β-cell SENP1 facilitates responsiveness to incretins and limits oral glucose intolerance in high fat fed mice

**DOI:** 10.1101/2020.11.30.402644

**Authors:** Haopeng Lin, Nancy Smith, Aliya F Spigelman, Kunimasa Suzuki, Mourad Ferdaoussi, Yaxing Jin, Austin Bautista, Ying Wayne Wang, Jocelyn E. Manning Fox, Jean Buteau, Patrick E MacDonald

**Author notes:** Correspondence to: Patrick MacDonald, Alberta Diabetes Institute, LKS Centre, Rm. 6-126, Edmonton, AB, Canada, T6G 2R3, email., web. www.bcell.org, twitter. @bcellorg.

## Abstract

SUMOylation reduces oxidative stress and preserves islet mass; but this happens at the expense of robust insulin secretion. To investigate a role for the deSUMOylating enzyme <u>sen</u>trin-specific <u>p</u>rotease <u>1</u> (SENP1) in glycemia following metabolic stress, we put pancreas/gut-specific SENP1 knockout mice (pSENP1-KO) on an 8-10-week high fat diet (HFD). Male pSENP1-KO mice were more glucose intolerant following HFD than littermate controls, but this was only obvious in response to oral glucose, and a similar but milder phenotype was observed in females. Plasma incretin responses were identical, and glucose-dependent insulinotropic polypeptide (GIP) was equally upregulated after HFD, in pSENP1-KO and - WT littermates. Islet mass was not different, but insulin secretion and β-cell exocytotic responses to Exendin4 (Ex4) and GIP were impaired in islets lacking SENP1. Glucagon secretion from pSENP1-KO islets was also reduced, consistent with the expected SENP1 knockout in all islet cells, so we generated β-cell–specific SENP1 knockout mice (βSENP1-KO). These phenocopied the pSENP1-KO mice with selective impairment in oral glucose tolerance following HFD, preserved islet mass expansion, and impaired β-cell exocytosis and insulin secretion to Ex4 and GIP. Thus, β-cell SENP1 limits glucose intolerance following HFD by ensuring a robust facilitation of insulin secretion by incretins such as GIP.

## Introduction

Glucose metabolism is the primary driver for insulin secretion, increasing the production of ATP, stimulating electrical activity and Ca^2+^ entry to trigger the exocytosis of insulin granules. Multiple additional factors serve either to maintain or augment the pool of secretory granules available to respond to the Ca^2+^ increase^1^. These pathways could be required for robust insulin secretory responses to receptor-mediated secretagogues such as the incretin hormones glucose-dependent insulinotropic polypeptide (GIP) and glucagon-like peptide-1 (GLP-1)^2^ which amplify glucose-stimulated responses via G-protein coupled receptors through cAMP-dependent and cAMP-independent mechanisms^3,4^. The action of the incretins to facilitate insulin secretion is impaired in type 2 diabetes (T2D), although the underlying mechanism for such a failure appears complex^5^ and could involve altered incretin receptor signalling^6^. Our recent study showed that incretin-induced insulin secretion from human islets correlates with the ability of glucose to augment depolarization-induced insulin exocytosis in single β-cells^7^. In transgenic mice with impaired metabolism-insulin granule coupling, achieved by β-cell specific deletion of the sentrin-specific protease-1 (SENP1), the β-cell response to the GLP-1 receptor agonist Exendin4 (Ex4) was impaired and the ability of DPP4 inhibition to improve oral glucose tolerance was greatly reduced^7^.

SENP1 couples upstream reducing equivalents from glucose metabolism to the downstream exocytotic site at the plasma membrane to facilitate exocytosis^8–11^. This redox-dependent pathway appears impaired in β-cells from human donors with type 2 diabetes (T2D)^12^, likely a result of upstream mitochondrial dysfunction. Loss of islet SENP1 results in moderate oral glucose intolerance with impaired insulin secretion, suggesting that deSUMOylation mediated by this enzyme is required for maintenance of glucose homeostasis^12^. Somewhat contradicting this we also found that overexpression of SENP1 induces apoptosis in β-cells^13^, consistent with the findings of others who showed the development of diabetes due to loss of islet mass in mice lacking the SUMO-conjugating enzyme Ubc9^14^. Up-regulation of islet Ubc9 protected from oxidative stress induced by streptozotocin (STZ), staving off overt diabetes, but impaired glucose-dependent insulin secretory responses. Effectively, SUMOylation appears required for the maintenance of β-cell viability at the cost of β-cell function^15^. It remains unknown whether the deSUMOylating enzyme SENP1 is required for the facilitation of insulin secretory responses and glucose tolerance under metabolic stress, such as high fat diet (HFD), or conversely whether loss of SENP1 would protect against glucose intolerance under these conditions by preserving β-cell mass and insulin secretion.

Here we investigated two interrelated questions in both gut/pancreas specific (pSENP1-KO) and β-cell specific (βSENP1-KO) SENP1 knockout mice. We asked whether the loss of islet/β-cell SENP1 sensitizes mice to HFD-induced glucose-intolerance; and we studied whether altered incretin responsiveness contributes to glucose intolerance under these conditions. We find that after an 8-10-week HFD, both pSENP1-KO mice and βSENP1-KO show a worsening of oral glucose intolerance compared with littermate controls. This is accompanied by a decreased insulin secretory response of β-cells to glucose, GLP-1 and GIP, without any difference in β-cell mass. Our findings support a model whereby SENP1 is required to ensure the availability of releasable insulin granules on which incretin signalling acts to augment glucose-stimulated insulin secretion. This becomes particularly important for limiting glucose intolerance following high fat feeding where incretin responses, notably plasma GIP responses, are increased.

## Methods

### Animals and diets

Pdx1-Cre mice (B6.FVB-Tg (Pdx1-cre)6 Tuv/J, Jackson Lab, 014647) on a C57BL/6 genetic background and Ins1-Cre mice on a mixed C57BL/6 and SV129 background^16^ were crossed with Senp1-floxed mice on a C57BL/6 background^12^ to generate gut/pancreas specific knockout mice (Pdx1-Cre^+^;Senp1^fl/fl^ - pSENP1-KO) and β-cell specific knockout mice (Ins1-Cre^+^;Senp1^fl/fl^ - βSENP1-KO) as described previously^7,12^. Pdx1-Cre^+^;Senp1^+/fl^ mice (pSENP1-HET) and Ins1-Cre^+^;Senp1^+/fl^ mice (βSENP1-HET) were used as heterozygotes, respectively. Pdx1-Cre^+^;Senp1^+/+^ mice (pSENP1-WT), and Ins1-Cre^+^;Senp1^+/+^ mice (βSENP1-WT) were used as littermate controls. Genotypes were confirmed from ear notches by using REDExtract-N-Amp Tissue PCR kit (Sigma-Aldrich) and primers as previously described^12^, and loss of SENP1 expression was confirmed by nested PCR and western blot in different tissues. At 12 weeks of age, mice were fed with HFD (60 % fat; Bio-Serv, CA89067-471) for 8-10 weeks.

### Western Blotting

SENP1 antibody-C12 (dilution 1:500, Santa Cruz sc-271360) and β-actin (dilution 1:500, Santa Cruz sc-47778) were used as primary antibodies. Mouse islets, gut and brain were homogenized with 7M Guanidine HCl using an IKA-WERK ULTRA-TURRAX homogenizer to extract proteins. The protein was precipitated by addition of methanol, chloroform and water in 4:1:3 ration (v/v). Protein pellet was recovered by centrifugation at 10,000 rpm for 10 minutes and then dissolved in 1%SDS, 0.2M Tris, 10mM DTT, pH6.5. The protein concentration was roughly estimated by the absorbance at 280nm. 50 μg, 10 μg and 10 μg protein from islets, intestine and brain respectively were loaded and separated by SDS-PAGE (7.5% gel), transferred to PVDF membrane, and then probed with the antibody in the presence of 5% skim milk.

### Nested qPCR of SENP1

Cre-lox recombination was used for the ablation of Exon14 and Exon15 of SENP1 gene in SENP1 KO mouse. To evaluate the efficiency of SENP1 KO, the cDNA encoding Exon14 and 15 was quantified by qPCR. Due to the low copy number of SENP1, the nested qPCR was used to increase the detection specificity and sensitivity of SENP1. Total RNA was extracted from kidney, brain, stomach, intestine and islets using TRIzol reagent (Fisher, 15596018). The cDNA was prepared from total RNA (50-100ng) with ABM 5x All-In-One RT Master Mix (G486, Applied Biological Materials Inc.). The cDNAs of SENP1 and cyclophilin A were amplified using preamplification primers and Platinum Taq DNA polymerase (10966-018, Invitrogen) with cycling parameter: 15 cycles of 94°C for 30 s, 60°C for 10s, 55°C for 10s and 72°C for 25 s. To remove Taq DNA polymerase and the primers, the PCR fragment was incubated with 4M Guanidine HCl for 10 minutes at room temperature and then precipitated with 80% ethanol in the presence of glycogen (50mg/ml). The nested qPCR was carried out with the qPCR primers, Fast SYBRGreen Master mix (438512, Applied Biosystems), 7900HT Fast real-time PCR system (Applied Biosystems) and the pre-amplified cDNAs as the templates with cycling parameter: 40 cycles of 95°C for 5 s, 50°C for 20s. All primers are listed in **Supplementary table 1**.

### In vivo studies

Mice on chow diet (CD) were fasted 4-5 hours prior to oral glucose tolerance test (OGTT) (1 g/kg dextrose)^17^ or intraperitoneal glucose tolerance tests (IPGTT) (1 g/kg dextrose)^18^ at 10-11 weeks. After 8 weeks of HFD mice were challenged with 0.5 g/kg dextrose in OGTT, and after 9 weeks of HFD mice were challenged with 1 g/kg dextrose in IPGTT after 9 weeks. Tail blood was collected for glucose and insulin measurement as previously described^12^. Insulin tolerance test (ITT)^19^ was performed on 12-week-old mice on CD or after 10 weeks HFD with IP injection of 1 U/kg Humulin R (Eli Lilly). To measure total plasma GLP-1 and GIP, after oral glucose gavage (2 g/kg dextrose), tail blood from mice on CD or following HFD was collected at indicated time and plasma was frozen until assay for total GLP-1 (Multi Species GLP-1 Total ELISA kit, Millipore) or GIP (Mouse GIP ELISA Kit, Crystal Chem).

### Pancreatic islet isolation and perifusion

Islets were isolated by collagenase digestion of the pancreas^20^ and cultured overnight. Insulin secretion was measured by perifusion as previously described^12^. Briefly, 25 islets were first pre-perifused for 30 minutes at 2.8 mM glucose before sample collection and treatments as with 16.7 mM glucose, Exendin-4 (Ex4, 100 nM, Sigma), GIP (100 nM, Anaspec), or KCl (30 mM) as outlined in the figures. Samples were collected every 1-5 minutes at a flow rate of 100 ul/min. At the end of the experiment, islets were lysed in acid/ethanol and all samples were stored at −20°C until assayed for insulin (STELLUX^®^ Chemi Rodent Insulin ELISA kit, Alpco). Glucagon perifusion solutions are described previously^21^. To assess the glucagon secretion, 85 islets were pre-perifused at 2.8 mM glucose for 30 min, followed sample collection and switching between 2.8 to 16.7 mM glucose, with GIP (100 nM, Anaspec) and alanine (10 mM, Sigma), as outlined in the figures. Samples were collected every 2 min at a flow rate of 100 ul/min. As with insulin, islets are lysed in acid/ethanol and stored at −20°C until assayed for glucagon (Rodent glucagon ELISA kit or U-Plex mouse glucagon ELISA kit, Mesoscale discovery).

### Patch-clamp electrophysiology

Islets were hand-picked and incubated with Ca^2+^-free buffer at 37°C for 10 minutes before being shaken and dispersed into single cells^9^. Patch-clamp was performed in the standard whole-cell configuration with sine+DC lockin function of EPC10 amplifier (HEKA Electronik). Extracellular and pipette solution was prepared as previously described.^22^ For Exendin-4 (Ex4, 100 nM, Sigma) and gastric inhibitory peptide (GIP,100 nM, Anaspec) stimulation, these were added to the extracellular solution with 5 mM glucose prior to the experiment. cAMP was not included in the pipette solution for these experiments. For glucose stimulation experiment^22^, cells were preincubated at 2.8 mM glucose for 1 hour and 0.1 mM cAMP was included in the pipette solution. Changes in capacitance were normalized to cell size (fF/pF). Mouse β-cells were identified by cell size (β-cell>4 pF) and half-maximal inactivation of Na^+^ currents at around −90 mV^23^.

### Histological analysis

Pancreata were weighed prior to being fixed in Z-fix (VWR Canada) and embedded in paraffin. Blocks were sectioned at 5 μm thickness with a total of 3-5 slides (each separated by 500 μm) being chosen for immunostaining and imaging as previously described^12^. Briefly, paraffin sections were rehydrated, permeabilized, blocked and incubated with guinea pig polyclonal insulin antibody (dilution 1:60, Dako, A0564) and mouse polycolonal glucagon antibody (dilution 1:1000, Sigma) overnight. Afterwards, sections were washed and incubated with Alex Fluor 488 (green) goat anti-guinea Pig IgG (dilution 1:500, A-11073-invitrogen) and Alexa Fluor 594 (red) goat anti-mouse IgG (dilution 1:500 A11037-invitrogen) for 1 hour. Last, they were washed and mounted in ProLong gold antifade reagent with DAPI (Life Technologies). To determine the islet area, insulin-positive cells were identified with tools from ImageJ (NIH) and normalized to total pancreas area. The islet mass was calculated as pancreas weight × relative islet area (as a proportion of pancreas section area). Islet size was determined by manual outlining in a blinded fashion using Zen Pro (Zeiss) and ImageJ (NIH).

### Statistical analysis

Graphpad Prism 8 for Mac OS X was used for student test, One-way or two-way ANOVA analysis followed by Bonferroni post-test to compare means between groups. Unbiased ROUT (robust regression followed by outlier identification) analysis was used for outlier identification and removal.

## Results

### Mice lacking islet SENP1 develop worse oral, but not intraperitoneal, glucose intolerance after HFD

Previously we showed that SENP1 was required for insulin granule exocytosis and that male pSENP1-KO mice had mild, but significant, oral glucose intolerance^12^. In the present study, we re-made our pSENP1-KO colony (the previous one lost Cre expression) by crossing new Pdx1-Cre^+^ mice from Jackson Labs (B6.FVB-Tg (Pdx1-cre)6 Tuv/J) and Pdx1-Cre^-^ Senp1^fl/fl^. We confirmed that β-cells from pSENP1-KO mice have impaired glucose-dependent facilitation of exocytosis (**Fig 1A**) as we’ve reported previously^12^ and we confirmed the loss of SENP1 in islets, along with a reduced expression in intestine and brain (**Fig 1B**). The pSENP1-KO mice have modest fasting hyperinsulinemia compared with littermate controls, but no differences in fasting glucose, body weight, or insulin tolerance (**Suppl Fig 1**). We performed OGTT, IPGTT, and ITT on both male and female mice at 10-12 weeks of age (**Fig 1C**). Male pSENP1-KO mice from this new colony were not obviously intolerant to oral or IP glucose (**Fig 1D, E; Suppl Figure 1A-D**), despite a reduced glucose-stimulated plasma insulin (**Fig 1F**). Female mice exhibited a similar phenotype, with some indication of IP glucose intolerance (**Fig 1G-I; Suppl Figure 1E-H**).

**Figure 1.**
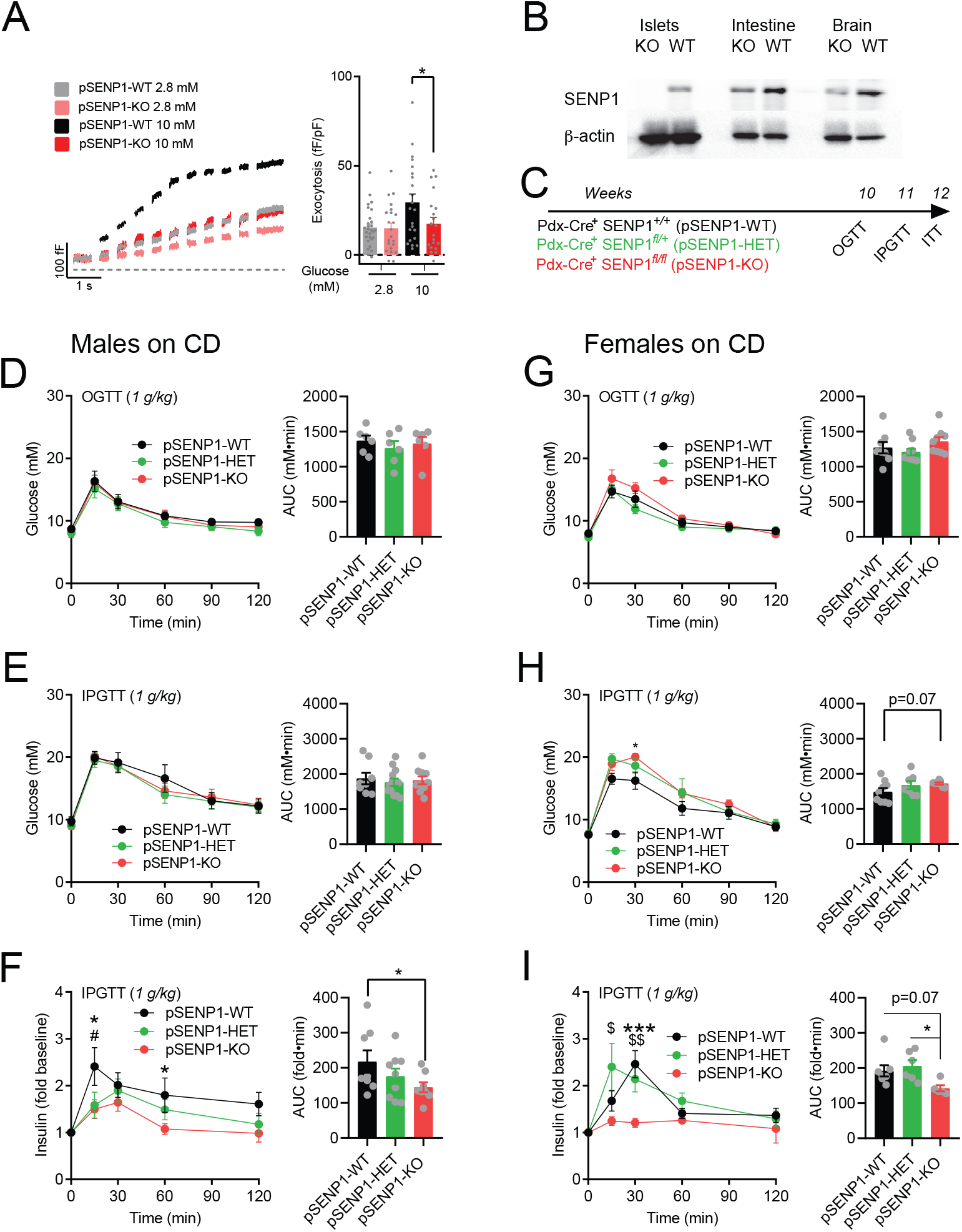
Normal glucose tolerance, but impaired insulin responses, of pSENP1-KO mice. **(A)** Representative traces, and average total responses, of β-cell exocytosis elicited by a series of membrane depolarizations at 2.8 and 10 mM glucose (n=28, 22, 22, 20 cells, n values correspond to graph bars from left to right, respectively). **(B)** Western blot of SENP1 expression in tissues from pSENP1-WT and pSENP1-KO mice. **(C)** schematic of experiments on chow diet (CD) fed mice. **(D)** Oral glucose tolerance test (OGTT) of male pSENP1-WT, -HET, and -KO mice (n=6, 6, 6 mice). **(E, F)** Intraperitoneal glucose tolerance (IPGTT) in male pSENP1-WT, -HET, and -KO mice (E - n=8, 13, 10 mice) and associated plasma insulin responses (F - n=8, 9, 8 mice). **(G)** OGTT of female pSENP1-WT, -HET, and -KO mice (n=8, 9, 9 mice). **(H, I)** IPGTT in female pSENP1-WT, -HET, and -KO mice (H - n=9, 7, 7), and associated plasma insulin responses (I - n=8, 6, 5 mice). AUC - area under the curve. Data are mean ± SEM and were compared with one-way or two-way ANOVA followed by Bonferroni post-test. *-p<0.05, **-p<0.01, ***-p<0.001 compared with the pSENP1-WT, while #-p<.05 indicated comparison between pSENP1-HET and pSENP1-WT. $-p<0.05, $$-p< 0.01 indicated comparison between pSENP1-HET and pSENP1-KO.

Together, these results confirm that SENP1 in the islets is important for insulin secretion, but this has modest, if any, impact on glucose tolerance under these conditions.

To further investigate the role of SENP1 in glucose homeostasis under high metabolic demand which mimics the progression towards T2D, we put mice on HFD for 8-10 weeks and assessed glucose homeostasis (**Fig 2A**). Without an obvious difference in insulin tolerance or fasting insulin (**Fig 2B, C**), male pSENP1-KO mice exhibited slightly higher fasting glucose and body weight compared to pSENP1-WT littermates (**Fig 2D, E**). In addition, male pSENP1-KO mice were modestly, but not significantly, intolerant of IP glucose (0.5 g/kg) compared with pSENP1-WT littermates (**Fig 2F**). They were however much more intolerant of an oral glucose challenge (0.5 g/kg) compared with littermate controls, with decreased plasma insulin responses (**Fig 2G, H**). Surprisingly, when we bypassed the gut to administer a higher dose of IP glucose (1 g/kg), there was still no obviously worse glucose intolerance in the pSENP1-KO mice (**Fig 2I**). In contrast, glucose intolerance to a higher oral glucose challenge (1 g/kg) was much more severely impaired with reduced glucose-stimulated plasma insulin secretion (**Fig 2J, K**), suggesting an important role for an impaired incretin effect in the pSENP1-KO mice. In our hands, and similar to what others have reported^24^, the female mice were generally resistant to developing insulin and glucose intolerance following HFD. Because of this, we did not observe any worsening of oral glucose intolerance in the female pSENP1-KO mice (**Suppl Fig 2A-F**). Taken together SENP1 deletion in the pancreas, and perhaps the gut and brain, impairs glucose stimulated insulin secretion and worsens oral, but not IP, glucose tolerance in male pSENP1-KO mice following HFD.

**Figure 2.**
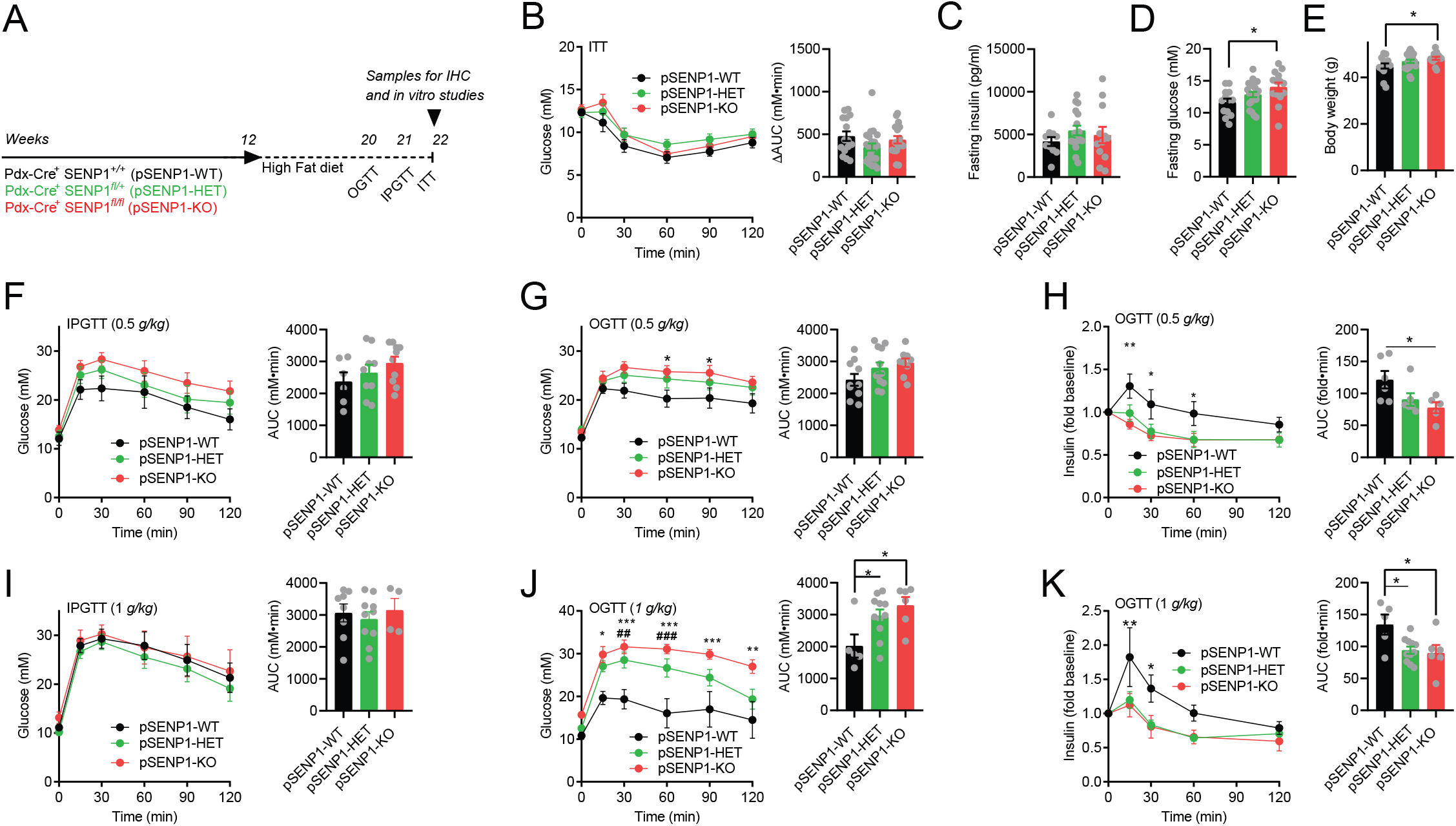
Selective worsening of oral glucose tolerance in pSENP1-KO mice after high fat feeding. **(A)** Schematic diagram of experiment on HFD. **(B)** Insulin tolerance test (ITT), **(C)** fasting insulin, **(D)** fasting glucose and **(E)** body weight of male pSENP1-WT, -HET, -KO mice on HFD (B - n=15, 22, 17 mice; C - n=10, 17, 12 mice; D - n=13, 21, 14 mice; E - =14, 23, 14 mice). **(F)** IPGTT with 0.5 g/kg of dextrose in male pSENP1-WT, -HET, and -KO mice after HFD (n= 6, 9, 10 mice). **(G, H)** OGTT (0.5 g/kg dextrose) of male pSENP1-WT, -HET, and -KO mice after HFD (G - n=10, 12, 9 mice), and associated plasma insulin responses (H - n=6, 6, 5 mice). **(I)** IPGTT with 1 g/kg of dextrose in male pSENP1-WT, - HET, and -KO mice after HFD (n=8, 10, 4 mice). **(J, K)** OGTT (1 g/kg dextrose) of male pSENP1-WT, - HET, and -KO mice after HFD (J - n=5, 11, 6 mice), and associated plasma insulin responses (K - n=5, 11, 6 mice). AUC - area under the curve. Data are mean ± SEM and were compared with student t test, one-way or two-way ANOVA followed by Bonferroni post-test. *-P < 0.05, **-P < 0.01, ***-P < 0.001 compared with the pSENP1-WT.

### β-cells from HFD-fed pSENP1-KO exhibit defective responses to glucose and incretins

A previous study showed that increasing SUMOylation protects islet mass from oxidative stress and apoptosis induced by the chemical agent streptozotocin^14^, and our own work suggests that knockdown of SENP1 protects β-cells from apoptosis^13^. To assess if islet SENP1 impacts β-cell mass in response to metabolic stress, we compared the islet mass between pSENP1-WT and pSENP1-KO mice following HFD. We found no significant differences in β-cell mass, islet number, and islet size between pSENP1-KO and pSENP1-WT littermates after HFD either in males (**Fig 3A**) or females (**Suppl Fig 3**), suggesting that SENP1 plays little or no role in β-cell mass under these conditions.

**Figure 3.**
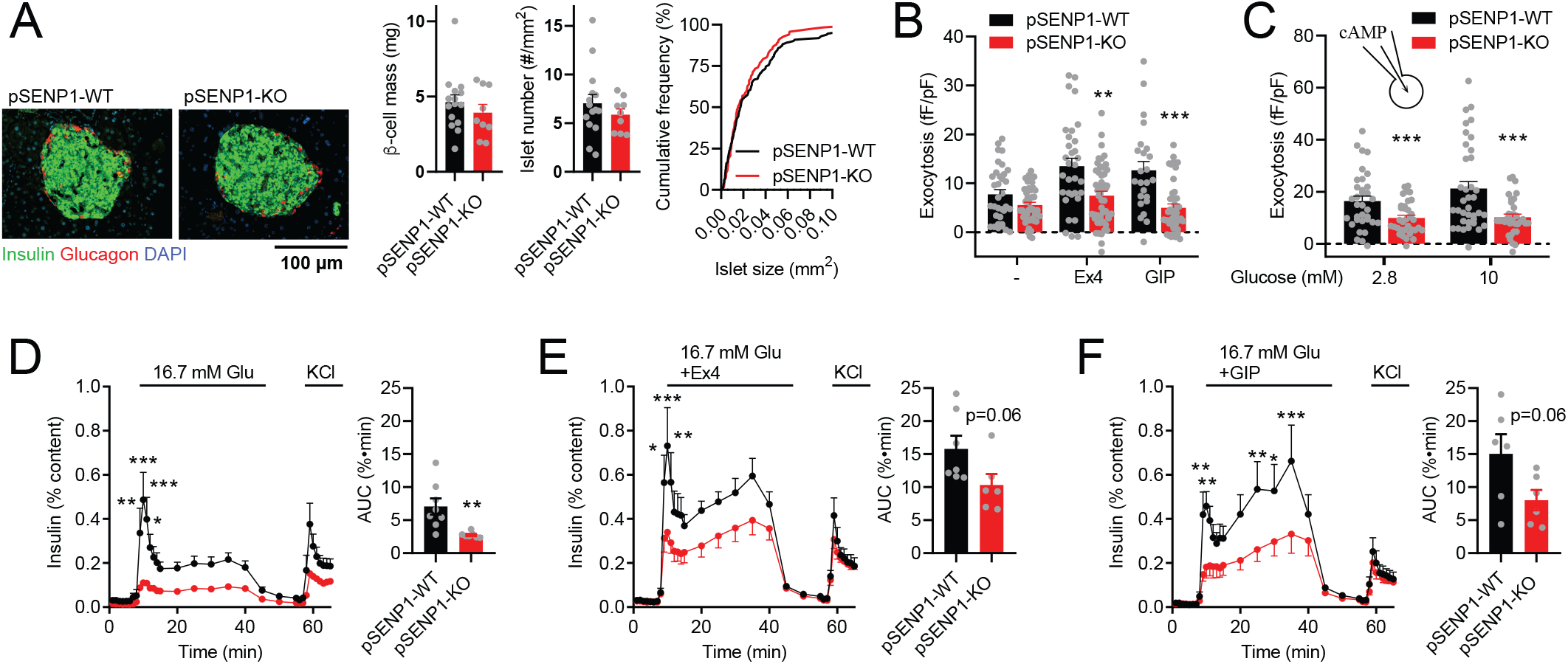
Impaired insulin secretion and exocytosis to glucose and incretins from male pSENP1-KO mice following HFD. **(A)** Representative immunostaining and quantification of β-cell mass, islet number, and islet size distribution in male pSENP1-WT (n =5 mice, 15 sections and 220 islets) and pSENP1-KO (n =3 mice, 9 sections and 116 islets) pancreas following HFD. Insulin (green), glucagon (red), and nuclei (blue). Scale bar=100 μm. **(B)** β-cell exocytosis, following HFD, in the presence of 5 mM glucose alone, and together with Exendin-4 (Ex4, 100 nM) or GIP (100 nM) (n=24-42 cells from 3-5 pairs of male mice). **(C)** β-cell exocytosis, following HFD, at 2.8 mM and 10 mM glucose with 0.1 mM cAMP included in pipette solution (n=34-37 cells from 3-4 pairs of male mice). **(D-F)** Insulin secretion from male pSENP1-WT and -KO islets following HFD in response to glucose alone (D - n=7, 6), or together with Ex4 (100 nM) (E - n=7, 6) or GIP (100 nM) (F - n=6, 6). AUC - area under the curve. Data are mean ± SEM and were compared with student t test, one-way or two-way ANOVA followed by Bonferroni post-test. *-p<0.05, **-p<0.01, ***-p<0.001 compared with the pSENP1-WT.

Since only OGTT, but not IPGTT, was impaired in pSENP1-KO mice after HFD, this prompted us to examine a role for incretin signaling in islets of these mice. To this end, single β-cells from pSENP1-KO mice fed HFD showed lower exocytotic responses, and this was much more obvious in the presence of the GLP-1R agonist Exendin-4 (Ex4, 100 nM) or GIP (100 nM; **Fig 3B**), suggesting that β-cell SENP1 is required for incretin-dependent amplification of insulin exocytosis. Previous reports suggested that SUMOylation inhibits GLP-1R activity and cell surface expression^25,26^. If reduced incretin receptor activity is responsible for the impaired Ex4/GIP-facilitated exocytosis we observe in the pSENP1-KO β-cells, then we should be able to rescue this effect by direct infusion of cAMP into these cells as we have previously shown in double-incretin receptor knockout (DIRKO) β-cells^27^. However, even with direct infusion of cAMP, the exocytotic response of β-cells from pSENP1-KO mice following HFD was still much lower than in β-cells from littermate controls (**Fig 3C**). This indicates that SENP1 is required for β-cell exocytosis downstream of cAMP signaling. Accordingly, we find that insulin secretion in response to glucose, Ex4 and GIP was lower in islets from HFD-fed pSENP1-KO mice (**Fig 3D-F**) vs control littermates, indicative of impaired incretin responsiveness in pSENP1-KO mouse islets.

### β-cell specific deletion of SENP1 leads to impaired OGTT after HFD

With the Pdx1 promotor as the driver of Cre expression, we expect loss of SENP1 will not be restricted to only β-cells **(Fig 4A**)^28^. Indeed, qPCR data showed that *Senp1* was around 80% lost in the proximal intestine and 100% absent from islets of pSENP1-KO mice (**Fig 4B**). This suggests that impaired oral glucose tolerance could be due to deletion of SENP1 from incretin-producing intestinal cells or glucagon-producing α-cells, rather than the loss of SENP1 in β-cells *per se*. However, total GLP-1 and GIP secretion *in vivo* were similar between pSENP1-WT and pSENP1-KO mice either on CD or HFD (**Fig 4C, D**). Our previous work showed that knockdown of SENP1 in α-cells can affect glucagon secretion^21^, and it is possible that this could impact insulin secretion incirectly^27^. Moreover GIP, which is upregulated in the HFD mice, is a potent stimulator for glucagon secretion^29^ as is the amino acid alanine^30^. We find here that glucagon secretion from pSENP1-KO islets in response to GIP and alanine is impaired in comparison with pSENP1-WT littermates (**Fig 4E**). Similar results were observed from islets of pSENP1-KO mice following HFD (not shown).

**Figure 4.**
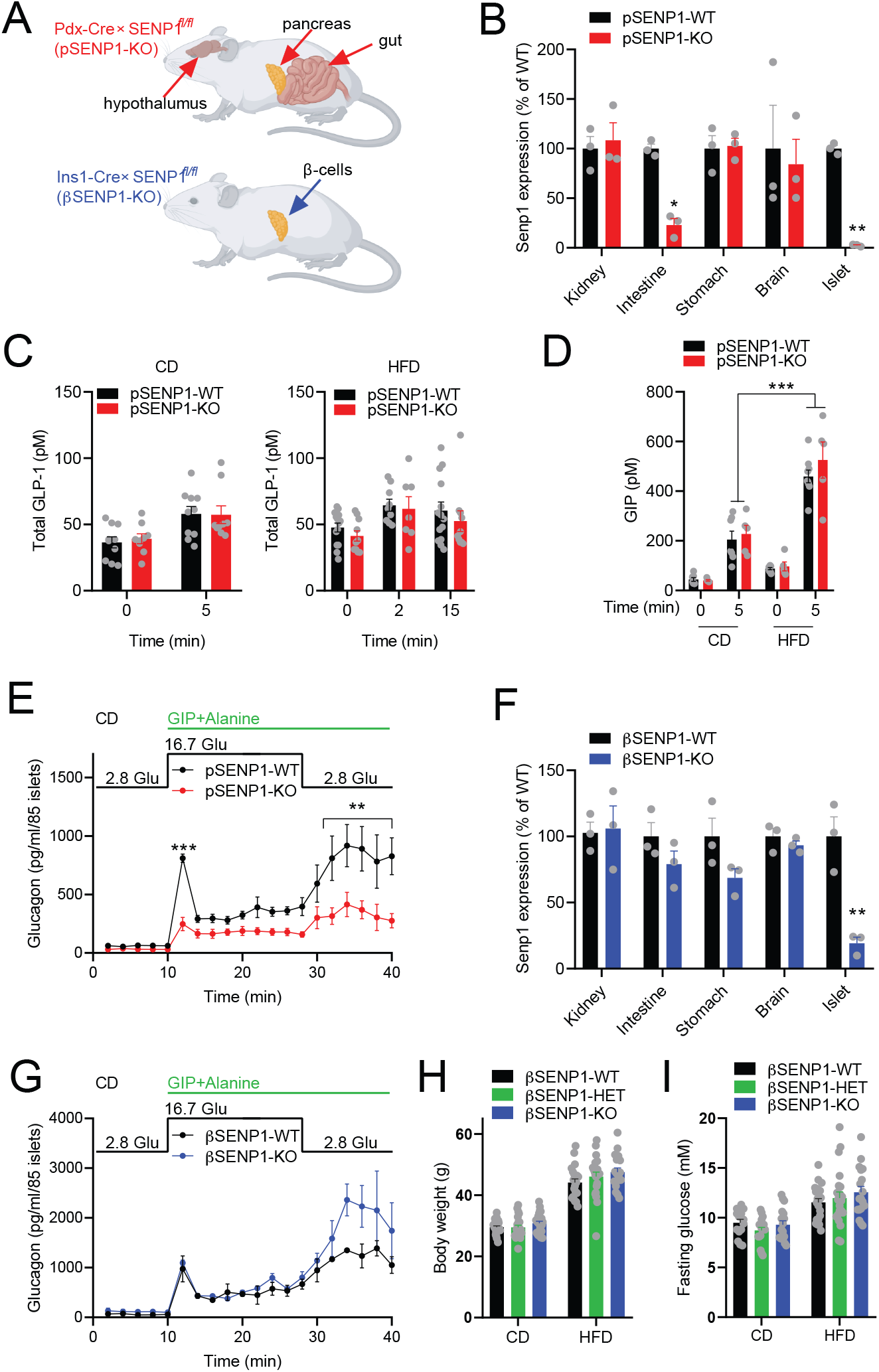
Generation of a β-cell specific SENP1 knockout. **(A)** Expected tissue selectivity of SENP1 knockout in the pSENP1-KO and βSENP1-KO mice. **(B)** qPCR of *Senp1* expression in tissues from pSENP1-WT and -KO mice (n=3, 3). **(C)** Oral glucose-stimulated total plasma GLP-1 in chow fed (CD, n=10, 9) and HFD mice (n=10, 9). Data from male and female were combined. **(D)** Oral glucose-stimulated plasma GIP in CD mice (n=7, 5) and HFD mice (n=9, 5). Data from male and female were combined. **(E)** Glucagon secretion at indicated glucose level in the presence of GIP and alanine in islets from pSENP1-WT and -KO mice on CD (n=3, 3). Data from male and female islets were combined. **(F)** qPCR of *Senp1* expression in tissues from βSENP1-WT and -KO mice (n=3, 3). **(G)** Glucagon secretion from islets of βSENP1-WT and -KO mice on CD (n=2, 2). Data from male and female islets were combined. **(H)** Body weight (n=18-23 mice) and **(I)** fasting glucose (n=15-22 mice) levels in male βSENP1-WT and -KO mice on CD and following HFD. Data are mean ± SEM and were compared with student t test, one-way or two-way ANOVA followed by Bonferroni post-test. *-p<0.05, **-p<0.01 compared with the pSENP1-WT.

Reduced glucagon responses may indirectly lower insulin secretion^27^. To differentiate the effect of β-cell from α-cell SENP1 on glucose homeostasis, we generated β-cell specific SENP1 knockout mice (βSENP1-KO) by crossing SENP1^fl/fl^ mice with the Ins1-Cre knock-in line^16^ (**Fig 4A**). qPCR confirmed that around 80% *Senp1* was lost in islets from βSENP1-KO, but was unaffected in other tissues (**Fig 4F**). Glucagon secretion from βSENP-KO islets was not different from that of βSENP1-WT littermates (**Fig 4G**), suggesting no effect on α-cell function in this model. We observed no significant differences in body weight, fasting glucose, insulin tolerance, or fasting insulin in these mice either on CD or following HFD (**Fig 4H, I; Fig 5A, B; Suppl Fig 4,5**). Similar to the pSENP1-KO mice, both oral and glucose tolerance were largely similar in βSENP1-KO and wild-type littermates at 8-10 weeks on CD in both males (**Fig 5C, D**) and females (**Suppl Fig 5D, E**). After 8-10 weeks of high fat feeding, similar to what we observed in the pSENP1-KO mice, male βSENP1-KO mice were more glucose intolerant than βSENP1-WT littermates to oral (**Fig 5E**) but not IP glucose (**Fig 5G**). Plasma insulin responses to oral glucose were impaired (**Fig 5F**), but this was not observed in response to IP glucose (**Fig 5H**). Similar, but less striking, differences were observed in the female pSENP1-KO mice (**Suppl Fig 5G-K**). Taken together, βSENP1-KO mice phenocopied the pSENP1-KO mice with an impaired insulin response and a worsened oral, but not IP, glucose tolerance following HFD; pointing to a β-cell specific SENP1-dependent incretin effect on glucose-homeostasis.

**Figure 5.**
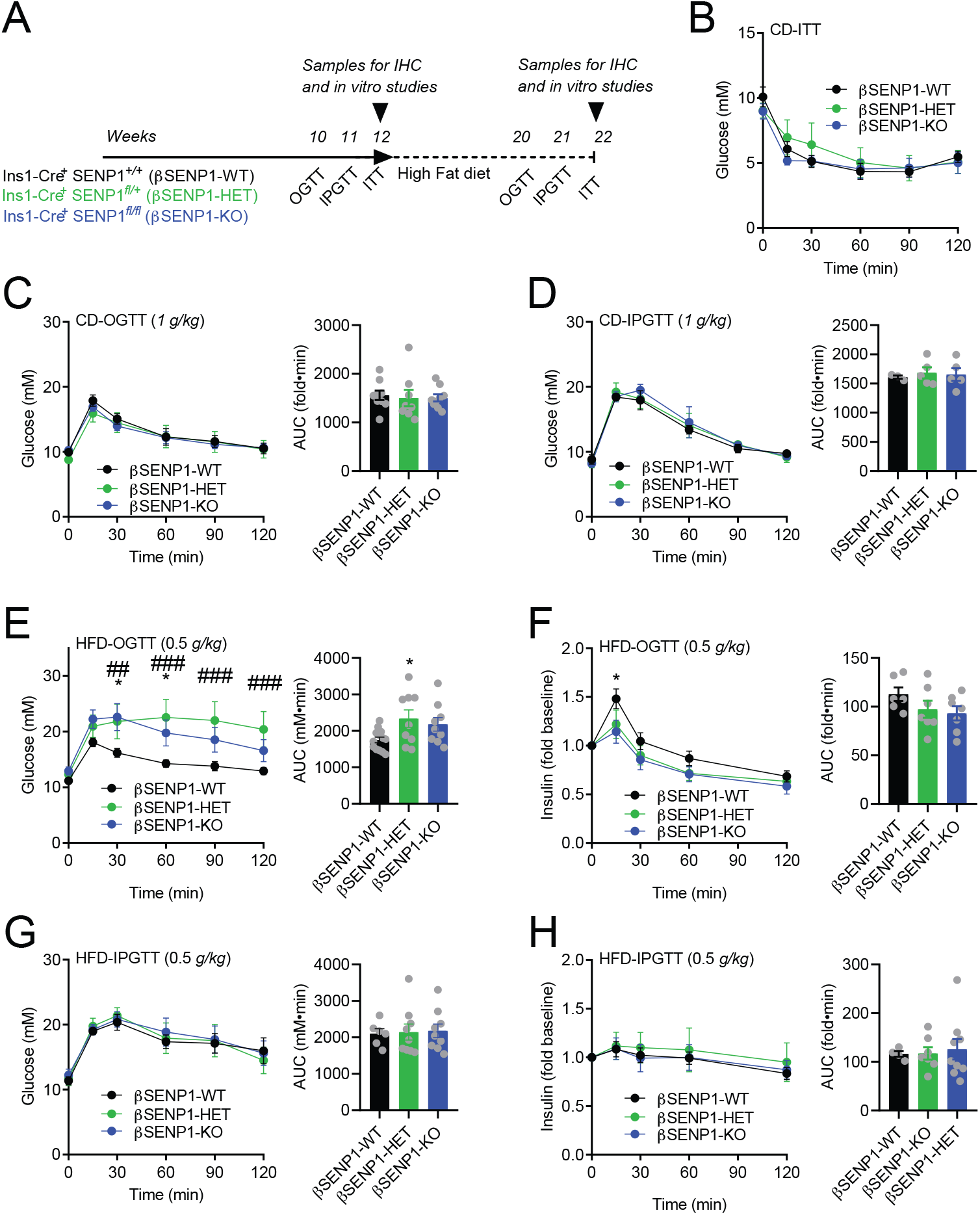
β-cell SENP1 knockout worsens oral, but not IP, glucose tolerance following HFD. **(A)** Schematic diagram of experiments on CD and HFD. **(B)** ITT of male βSENP1-WT, -HET, and -KO mice on CD (n=6, 6, 6 mice). **(C)** OGTT of male βSENP1-WT, -HET, and -KO mice on CD (n=9, 8, 9 mice). **(D)** IPGTT of male βSENP1-WT, - HET, and -KO mice on CD (n=3, 5, 5 mice). **(E, F)** OGTT of male βSENP1-WT, - HET, and -KO mice following HFD (E - n=13, 9, 9 mice), and associated plasma insulin responses (F - n=6, 7, 7 mice). **(G, H)** IPGTT (0.5 g/kg) of male βSENP1-WT, -HET, and -KO mice after HFD (G - n=6, 9, 9 mice), and associated plasma insulin responses (H - n=4, 9, 7 mice). AUC - area under the curve. Data are mean ± SEM and were compared with one-way or two-way ANOVA followed by Bonferroni post-test. *-p-<0.05 compared with the pSENP1-WT, while ##-p<0.01, ###-p<0.001 indicated comparison between pSENP1-HET and pSENP1-WT.

Similar to our observation in the pSENP1-KO model, we find no difference in β-cell mass, islet number, or islet size in βSENP1-KO mice compared with littermate controls (**Fig 6A, B; Suppl Fig 6**). Islet mass expansion upon high fat feeding in this model is preserved. We confirm impaired single β-cell exocytosis (**Fig 6C**) and reduced glucose- and KCl-stimulated insulin secretion (**Fig 6D**) from isolated cells and islets of chow-fed βSENP1-KO mice. After HFD, insulin secretion from βSENP1-WT islets was impaired to a similar level as that observed in βSENP1-KO islets (**Fig 6E, H**). Insulin secretion from βSENP1-WT islets was, however, rescued to a much greater degree by Ex4 and GIP than observed in βSENP1-KO islets, as demonstrated on both first phase and second phase insulin secretion (**Fig6 F-H**). Finally, Ex4 and GIP were unable to increase exocytosis to the same extent from β-cells of HFD βSENP1-KO mice compared with littermate controls, and these differences were much larger than with glucose alone (**Fig 6I**). Again, exocytosis could not be rescued by direct infusion of cAMP (**Fig 6J**) which revealed a larger difference between the βSENP1-WT and -KO β-cells than that observed in the absence of cAMP. Taken together, these data suggest that SENP1 ensures the availability of a pool of granules on which incretin receptor signaling can act to facilitate insulin secretion, which becomes particularly important after HFD to limit oral glucose intolerance.

**Figure 6.**
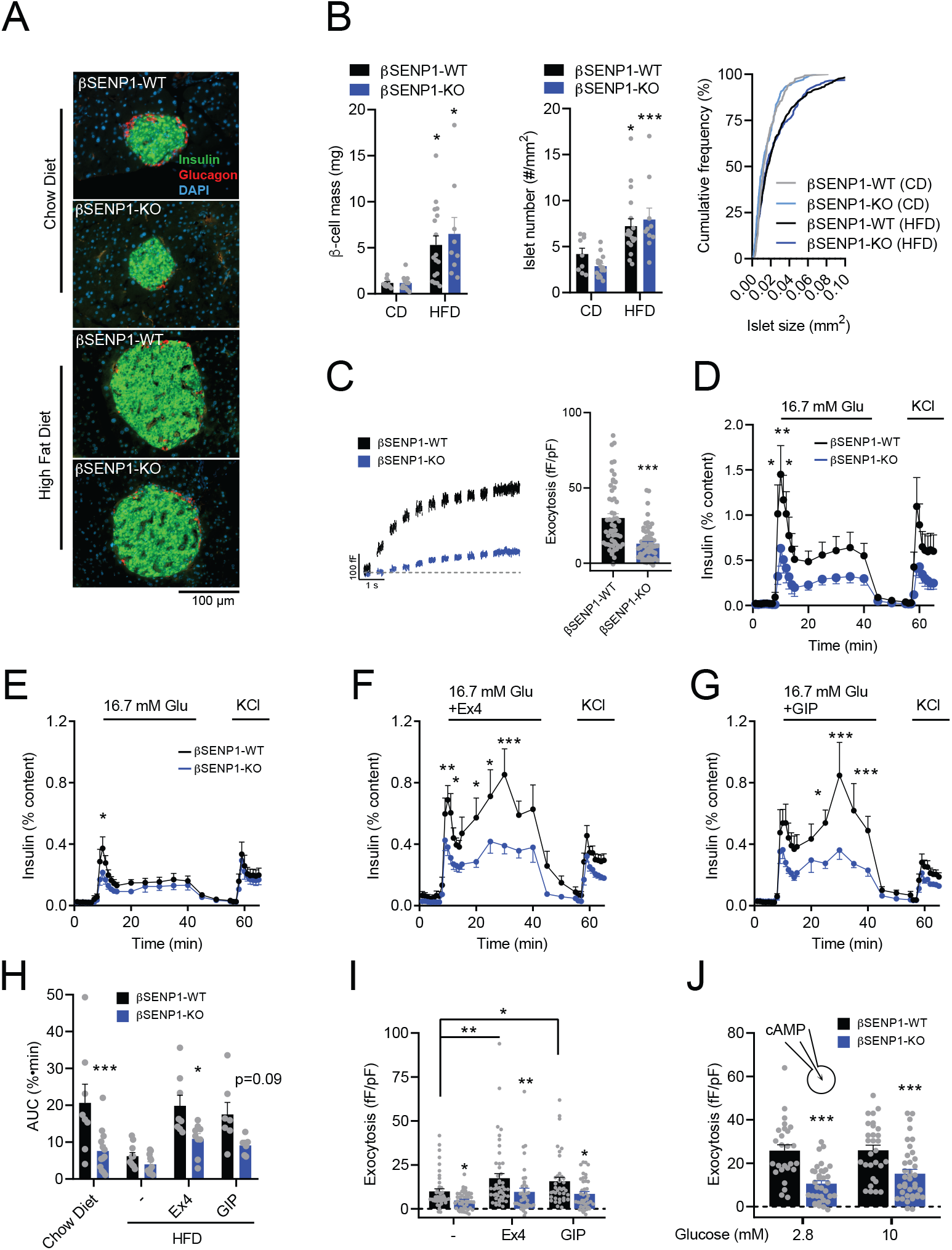
Impaired insulin secretion and exocytosis to glucose and incretins from βSENP1-KO islets following HFD. **(A)** Representative immunostaining images, and **(B)** β-cell mass, islet number, and islet size distribution of male βSENP1-WT mice on CD (n = 3 mice, 9 sections and 162 islets), male βSENP1-KO mice on CD (n = 4 mice, 13 sections and 139 islets), male βSENP1-WT mice following HFD (n = 6 mice,18 sections and 340 islets) and βSENP1-KO mice following HFD (n = 3 mice, 9 sections and 200 islets). Insulin (green), glucagon (red), and nuclei (blue). Scale bar=100 μm. **(C)** Exocytosis in single β-cells from male βSENP1-WT and -KO mice on CD at 10 mM glucose (n=55, 61 cells, n=5, 7 mice). **(D-G)** Insulin secretion from male βSENP1-WT and -KO islets from mice on CD in response to glucose (D - n=8, 7), or from male mice following HFD in response to glucose alone (E - n=9, 12) or together with Exendin-4 (Ex4, 100 nM; F - n=8, 9) or GIP (100 nM; G - n=7, 7). **(H)** Area under the curve (AUC) of the responses from panels D-G. **(I)** Exocytosis in β-cells of male βSENP1-WT and -KO mice following HFD at 5 mM glucose alone, or with Ex4 (100 nM) or GIP (100 nM) (n=44, 46, 47, 38, 40, 41 cells from 5 mice per group). **(J)** Exocytosis in β-cells of male βSENP1-WT and -KO mice following HFD at 2.8 mM and 10 mM glucose with 0.1 mM cAMP included in pipette solution (n=26, 35, 29, 38 cells from 3, 4, 3, 4 mice). Data are mean ± SEM and were compared with student t test, one-way or two-way ANOVA followed by Bonferroni post-test. *-p<0.05, **-p<0.01, ***-p<0.001 compared with the βSENP1-WT.

## Discussion

Here we aimed to investigate the role of the deSUMOylating enzyme SENP1 within the β-cell in glucose homeostasis following HFD-induced metabolic stress. Using pSENP1-KO and βSENP1-KO mouse models, we confirmed that loss of SENP1 in β-cells results in impaired exocytosis and reduced insulin secretion. This is consistent with the SUMOylation-dependent inhibition of insulin secretion shown by us and others insulinoma cells^11^, human β-cells^9,10,12^ and mouse β-cells^9,12,14^. It is clear that deSUMOylation, likely of multiple targets involved in exocytosis^31^, is an important mechanism facilitating insulin secretion. *In vivo*, transgenic over-expression in β-cells of the SUMO-conjugating enzyme Ubc9 leads to more obvious IP glucose intolerance^14^ than what we reported in our original pSENP1-KO mice^12^ while in our current pSENP1-KO and βSENP1-KO mice we find only a trend towards glucose intolerance, if any. In these mice we see impaired *in vivo* insulin responses to glucose, but this may be compensated for in part by an increased fasting insulin (Suppl Fig 1) thus limiting glucose intolerance in the absence of metabolic stress. We should also recognize that other pathways facilitating insulin secretion could compensate for the loss of SENP1 *in vivo*, for example lipid-mediated signaling^32,33^, which may be minimized in *in vitro* experiments performed in minimal buffers without fatty acid. Nonetheless, high fat feeding revealed the importance SENP1-dependent insulin secretion in the maintenance of glucose homeostasis.

It was somewhat surprising to us that a worsening of HFD-induced glucose intolerance in both the pSENP1-KO and βSENP1-KO mice was observed selectively in response to oral, but not IP, administration of glucose. This suggests an important interaction with the incretin response, which is upregulated in HFD as shown by others^34–36^ and in our observation that GIP responses in particular are enhanced. We can rule out the possibility that impaired incretin production is responsible for the worsened glucose intolerance in the SENP1 knockout models. Neither GIP nor GLP-1 responses were altered in pSENP1-KO mice, despite loss of *Senp1* in the proximal intestine, and the βSENP1-KO mice have preserved *Senp1* expression in the intestine but still exhibit a worsening of oral glucose intolerance after high fat feeding. Recent work has highlighted the important interaction between incretin responses, glucagon secretion, and control of insulin release^37^. In particular, GIP and amino acid induced glucagon secretion may be important in supporting robust insulin responsiveness^27,38–40^. Loss of SENP1 in α-cells could indirectly impact insulin secretion, and although we previously showed SUMO-dependent increases in glucagon secretion stimulated by adrenaline^21^, here we find an impaired glucagon response to GIP and alanine in the pSENP1-KO mice. It is possible that this contributes to a somewhat more severe phenotype in the pSENP1-KO than the βSENP1-KO mice, as effects on hyperglycemia and impaired *in vivo* and *in vitro* insulin responses were more severe in the former. However insulin tolerance was normal in the pSENP1-KO mice in this and our previous study^12^, suggesting that the effect of α-cell SENP1 on counter-regulatory function at least under hypoglycemic conditions where the β-cell is inactive, is negligible. Finally, the observation that HFD-induced oral glucose intolerance persists in the βSENP-KO model suggests that β-cell SENP1 is primarily responsible for the impaired incretin effect following HFD.

Consistent with this, isolated β-cells and islets showed defective exocytosis and insulin secretion. From βSENP1-KO islets following HFD in particular, this was most notable when GIP or Ex4 was present. On HFD, the demand for compensatory insulin secretion became elevated, with which SENP1-KO islets might not be able to catch up. Plasma GIP responses were much higher after HFD, suggesting that the adaptive GIP input and incretin signaling might compensate for the dysfunctional response to glucose and become more important after HFD^34^. *In vitro*, impaired insulin secretion from the βSENP1-WT islets following HFD could be partially rescued by GIP or GLP-1, but this response was absent in the βSENP1-KO islets. This could explain why we only observed worsened oral but not IP glucose tolerance in the absence of β-cell SENP1. A previous study showed that SENP1 might decrease GLP-1-induced insulin secretion by regulating the deSUMOylation of GLP-1 receptor^25,26^. However, when we measured exocytosis with high cAMP included in pipette solution to bypass the GLP-1 and GIP receptors, β-cell from βSENP1-KO still showed a decreased exocytotic response. This suggests that the impaired response to incretin resides downstream of receptor signaling, since this same experiment was able to rescue function in double-incretin-receptor knockout β-cells^27^. Altogether our data suggest that a robust insulin secretory response to incretin receptor activation, up-regulated after HFD in part by increased plasma GIP responses to oral glucose, requires metabolic signaling mediated by SENP1. It is possible that SENP1 is required to ensure availability of insulin granules on which cAMP-dependent signals act, or these converge on some common targets, such as synaptotagmin VII^9,41^.

Finally, SUMOylation protects against STZ-induced β-cell apoptosis and diabetes *in vivo* in mice^14^ and knockdown of SENP1 reduces cell death induced by inflammatory cytokines in insulinoma cells and human β-cells *in vitro*^13^. While up-regulation of SUMOylation therefore appears to protect from frank diabetes by reducing β-cell death, this comes at the cost of allowing robust upregulation of insulin secretion in response to metabolic stressors. Although we did not observe any differences in β-cell mass in the various models we studied, which suggests that SENP1 is not required for HFD-induced β-cell proliferation, it is possible that we could see preservation of β-cell mass over much longer periods of high fat feeding where an effect on β-cell apoptosis may be revealed. Likewise, we find that incretin signaling is important in limiting glucose intolerance following the 8-week HFD and that SENP1 facilitates this, but we do not know whether SENP1 activity in β-cells is required in concert with an upregulation of the incretin response or whether SENP1, known to couple cellular redox state to insulin secretory capacity, plays other roles in the progression of glucose intolerance at early or intermediate time points. In conclusion, our study shows that after an 8-week HFD, β-cell–specific SENP1 is dispensable in islet mass increases, but is required for incretin-induced insulin secretion that contributes to maintenance of oral glucose tolerance.

---

**Supplementary table 1.**
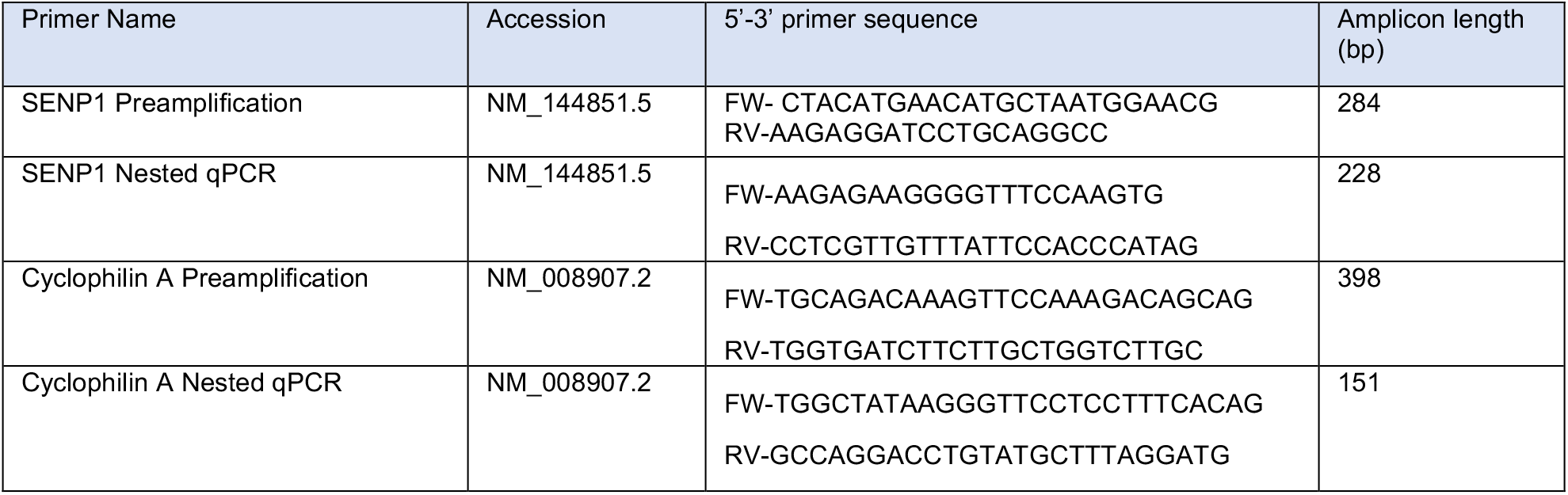

## Acknowledgements

The authors wish to thank Dr. Edward Yeh (University of Arkansas) who originally provided the SENP1^fl/fl^ mice, and Dr. Jon Campbell (Duke University) for many helpful discussions. This work was funded by a Foundation Grant to PEM from the Canadian Institutes of Health Research (CIHR: 148451). HL was supported by a from the Sino-Canadian Studentship from Shantou University. YJ was supported by a University of Alberta Office of the Provost and VP (Academic) Summer Studentship. PEM holds the Canada Research Chair in Islet Biology.

## Author Contributions

HL, NS, AS, MF, KS, YJ, YWW and AB researched and analyzed data. HL, JEMF, JB and PEM designed the studies. HL and PEM wrote the manuscript. All authors edited and approved of the final version. PEM acts as guarantor of the work.

