## Supplementary Figures for "β-cell SENP1 facilitates responsiveness to incretins and limits oral glucose intolerance in high fat fed mice"

**Supplementary Figure Legends**

**Supplementary Figure 1. Additional in vivo glucose homeostasis assessment of male and female pSENP1-KO mice on chow diet (CD).**

**(A)** Fasting insulin; **(B)** fasting glucose, **(C)** body weight, and **(D)** IP insulin tolerance of pSENP1-WT, -HET and -KO male mice on CD (n=6-22 mice). **(E)** Fasting insulin, **(F)** fasting glucose, **(G)** body weight, and **(H)** IP insulin tolerance of pSENP1-WT, -HET and -KO female mice on CD (n=7-17 mice).

AUC - area under the curve. Data are mean ± SEM and were compared with student t test, one-way or two-way ANOVA followed by Bonferroni post-test. *-p<0.05, **-p< 0.01, compared with the pSENP1-WT.

**Supplementary Figure 2. In vivo glucose homeostasis assessment of female pSENP1-KO mice following high fat diet (HFD).**

**(A)** OGTT and **(B)** associated plasma insulin responses, **(C)** IP insulin tolerance, **(D)** body weight, **(E)** fasting glucose, and **(F)** fasting insulin of pSENP1-WT, -HET and -KO female mice following HFD (n=9-14 mice).

**Supplementary Figure 3. Islet morphometry analysis of female pSENP1-KO mice after HFD.**

**(A)** Representative immunostaining, β-cell mass, islet number, and islet size distribution of female pSENP1-WT (n = 5 mice, 15 sections and 189 islets) and pSENP1-KO (n = 6 mice, 18 sections and 197 islets) following HFD. Insulin (green), glucagon (red), and nuclei (blue). Scale bar=100 µm. Data are presented as mean ± SEM.

**Supplementary Figure 4. Additional in vivo glucose homeostasis assessment of βSENP1-KO male mice.**

(A) IP insulin tolerance of male βSENP1-WT, -HET, and -KO mice following HFD (n=15, 8, 8 mice). (B) Delta area under the curve (ΔAUC) of IP insulin tolerance tests of male βSENP1-WT, -HET, and -KO mice on CD and following HFD (n=6, 6, 6, 15, 8, 8 mice). (C) Fasting insulin of male βSENP1-WT, -HET, and -KO mice on CD and following HFD (n=6, 4, 5, 13, 17, 14). Data are presented as mean ± SEM.

**Supplementary Figure 5. In vivo glucose homeostasis assessment of βSENP1-KO female mice.**

(A) Fasting insulin (n=6, 8, 4, 13, 16, 14 mice), (B) fasting glucose (n=21, 22, 20, 21, 21, 22), and (C) body weights (n=30, 23, 25, 20, 22, 22) of female βSENP1-WT, -HET, and -KO mice on CD and following HFD. (D) OGTT (n=11, 12, 12 mice), (E) IPGTT (n=10, 11, 9 mice), and (F) IP insulin tolerance (n=6, 9, 8) of female βSENP1-WT, -HET and -KO mice on CD. (G) OGTT (n=11, 13, 11 mice) and (H) associated plasma insulin response (n=7, 7, 8 mice) of female βSENP1-WT, -HET, and -KO mice following HFD. (I) IPGTT (8, 9, 10), and (J) associated plasma insulin response (n=6, 9, 6 mice) of female βSENP1-WT, -HET, and -KO mice following HFD. (K) IP insulin tolerance (n=11, 11, 12 mice) of female βSENP1-WT, -HET and -KO female mice following HFD and associated ΔAUC for CD and HFD. Data are presented as mean ± SEM.

**Supplementary Figure 6. Islet mass analysis of female βSENP1-KO mice after CD and HFD feeding.**

(A) Representative immunostaining image, (B) islet mass, islet number, islet size accumulative frequency of female βSENP1-WT mouse on CD (n = 5 mice, 14 sections and 146 islets ), βSENP1-KO on CD mouse (n = 4 mice, 12 sections and 101 islets), βSENP1-WT mouse following HFD (n = 5 mice,15 sections and 155 islets ) and βSENP1-KO mouse following HFD (n = 4 mice, 11 sections and 141 islets). Insulin (green), glucagon (red), and nuclei (blue). Scale bar=100 μm. Data are mean ± SEM and were compared with one-way or two-way ANOVA followed by Bonferroni post-test. *-P < 0.05 indicated comparison between HFD and CD.

### Supplementary Figure 1

Males on CD

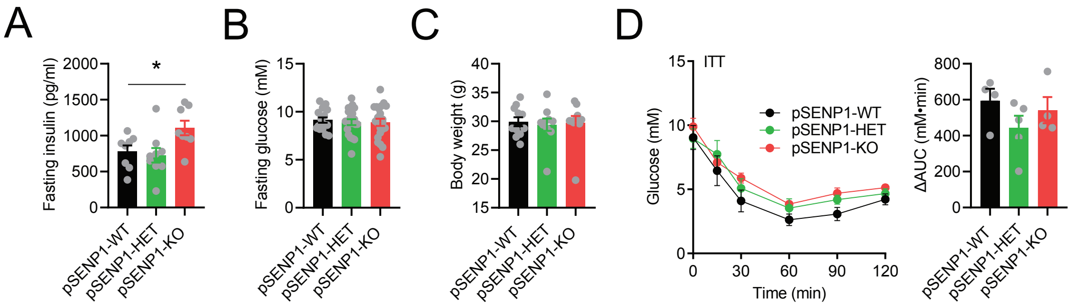


Females on CD

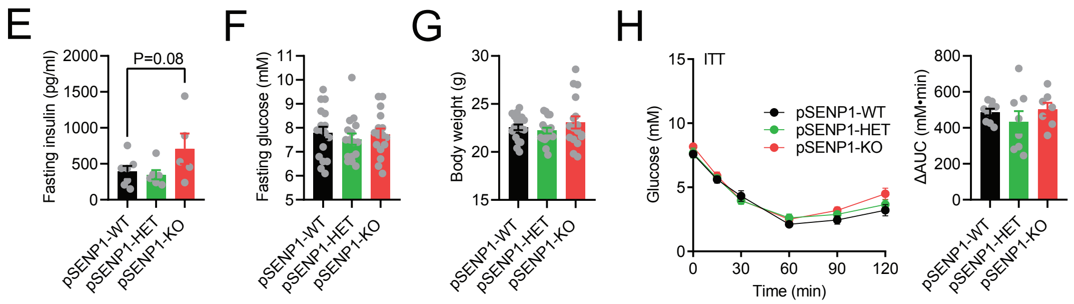

### Supplementary Figure 2

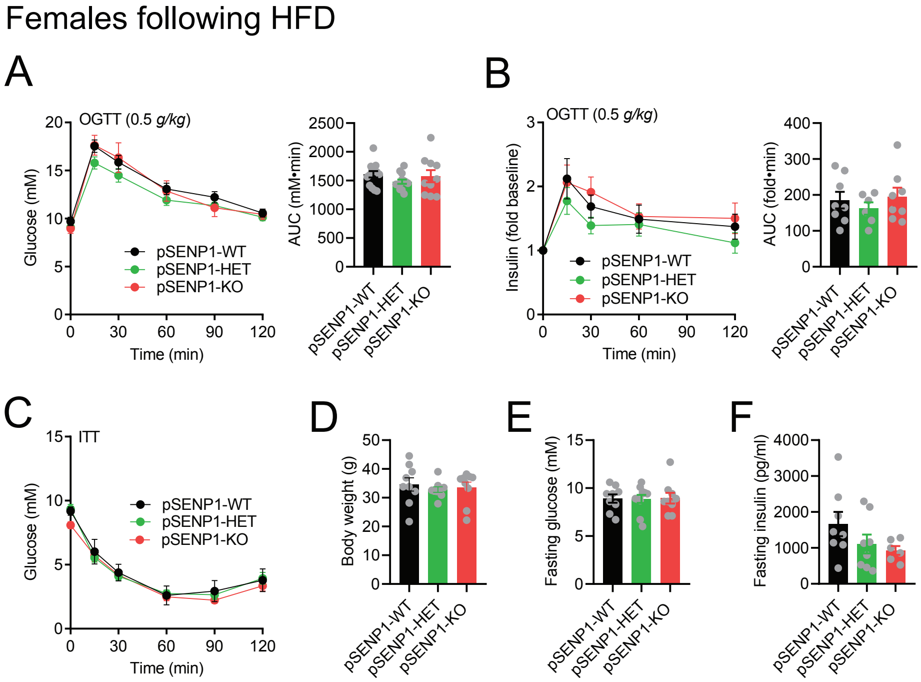

### Supplementary Figure 3

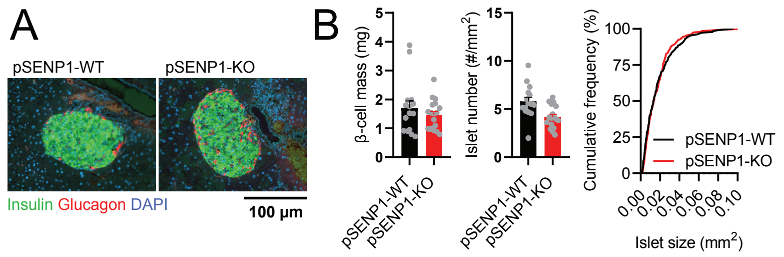

### Supplementary Figure 4

#### Male mice

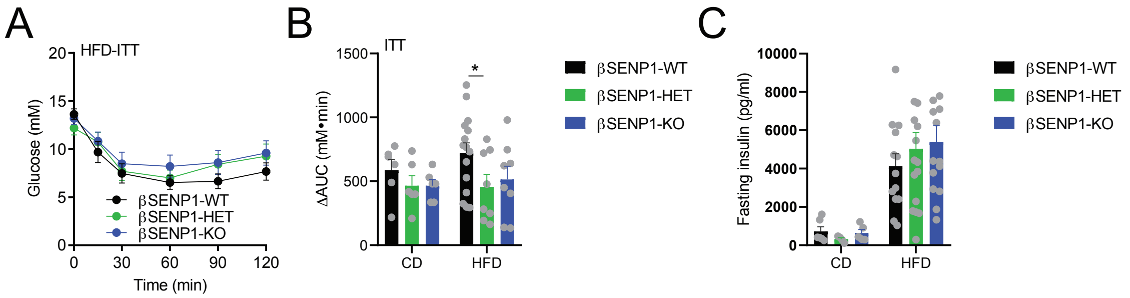

### Supplementary Figure 5

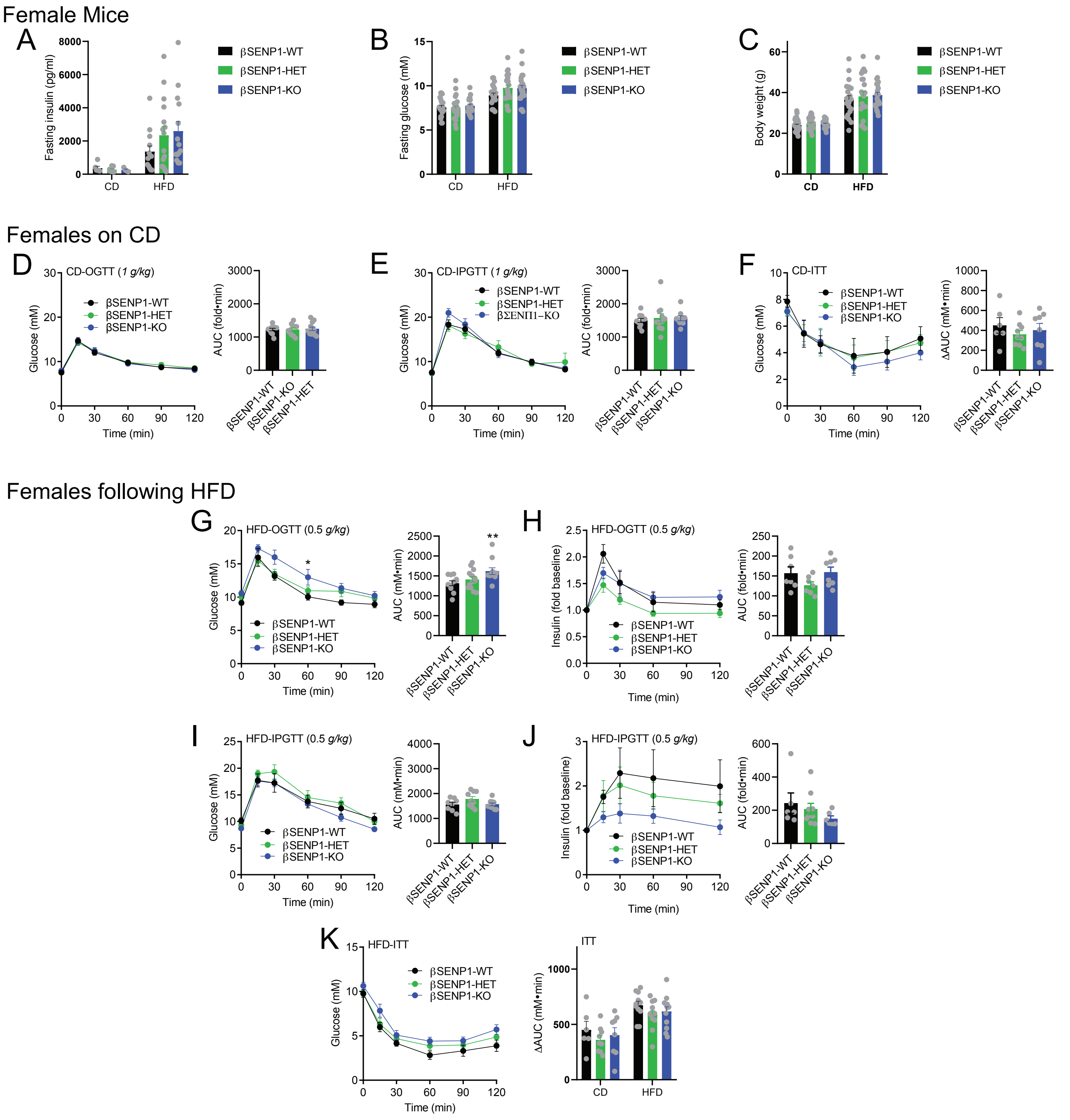

### Supplementary Figure 6

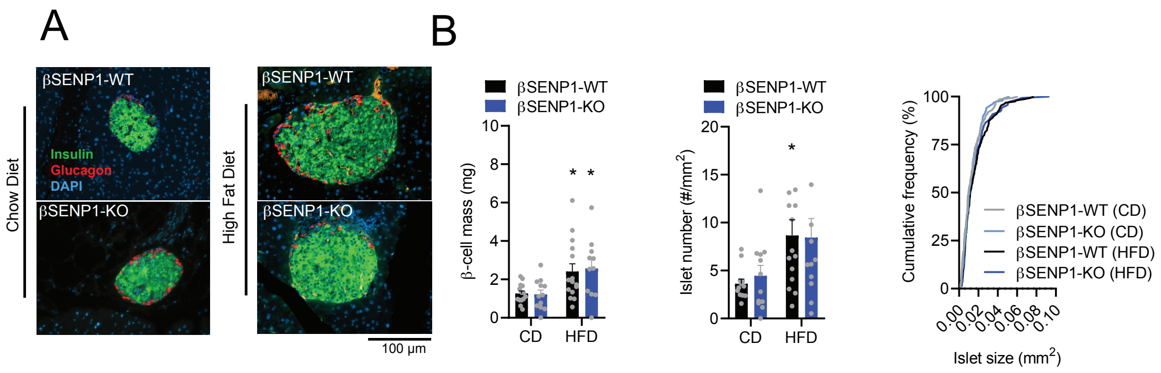
